# MRBLES 2.0: High-throughput generation of chemically functionalized spectrally and magnetically-encoded hydrogel beads using a simple single-layer microfluidic device

**DOI:** 10.1101/2020.06.22.166074

**Authors:** Yinnian Feng, Adam K. White, Jamin B. Hein, Eric A. Appel, Polly M. Fordyce

## Abstract

Widespread adoption of bead-based multiplexed bioassays requires the ability to easily synthesize encoded microspheres and conjugate analytes of interest to their surface. Here, we present a simple method (MRBLEs 2.0) for efficient high-throughput generation of microspheres with ratiometric barcode lanthanide encoding (MRBLEs) bearing functional groups for downstream bioconjugation. Bead production in MRBLEs 2.0 relies on manual mixing of lanthanide/polymer mixtures (each of which comprises a unique spectral code) followed by droplet generation using single-layer, parallel flow-focusing devices and off-chip batch polymerization of droplets into beads. To streamline downstream analyte coupling, MRBLEs 2.0 crosslinks copolymers bearing functional groups on the bead surface simultaneously during bead generation. Using the MRBLEs 2.0 pipeline, we generate monodisperse MRBLEs containing 48 distinct well-resolved spectral codes in high-throughput (>150,000/min and can be boosted to 450,000/min). We further demonstrate the efficient conjugation of oligonucleotides and entire proteins to carboxyl MRBLEs and biotin to amino MRBLEs. Finally, we show that MRBLEs can also be magnetized via simultaneous incorporation of magnetic nanoparticles with only a minor decrease in the potential code space. We anticipate that MRBLEs 2.0 can be directly applied towards a wide variety of downstream assays from basic biology to diagnostics and other translational research.

## Introduction

The recent ability to identify molecules across a cell’s genome, transcriptome, and proteome has dramatically increased the need for technologies capable of detecting interactions between biological macromolecules at scale to understand interactome networks. Multiplexed bioassays, in which binding is assessed between a single ‘bait’ molecule and many possible ‘prey’ interactors, can reduce the number of experiments required to explore a potential interactome space and speed the pace of discovery. Pioneering examples of these assays used spatial arrays to test thousands to millions of potential interactions in a single experiment^1,2^. However, spatial arrays suffer from relatively slow kinetics (as bait molecules are immobilized on a planar surface) and typically require relatively large amounts of sample^1^.

Multiplexed bead-based assays provide an appealing alternative, providing near fluid-phase interfacial kinetics, many replicates per experiment, opportunities for quality control, and the ability to flexibly couple different probes and targets across experiments^3-5^. Spectrally encoded beads, in which beads are embedded with ratiometric combinations of fluorescent or luminescent materials, provide a particularly convenient format for multiplexed assays and have already been used for a wide variety of applications^6-16^. To date, Luminex multi-analyte profiling (xMAP) technology represents the most widely used spectrally encoded bead-based technology. Luminex xMAP beads are commercially available and compatible with flow cytometry^17^ and are widely used for a variety of bioassays^18^. These beads consist of magnetic polystyrene microspheres that encapsulate distinct proportions of red and infrared fluorophores, each of which comprises a unique spectral code. However, these beads also suffer from a variety of limitations. First, the use of fluorescent dyes for encoding limits the possible coding space to 500 and code sets typically contain <100 codes. Second, hydrophobic polystyrene beads cannot detect low affinity interactions due to widespread nonspecific binding mediated by hydrophobic interactions^19^.

Recently developed spectrally encoded hydrogel beads provide an appealing alternative that have been used for a variety of biomedical and sensing applications^20,21^. Hydrogel beads are comprised of a cross-linked, hydrated polymeric network made up of one or more hydrophilic monomers, providing near fluid-phase kinetics at functionalized surfaces as well as high-efficiency molecular loading^22,23^, and microfluidic droplet generators have been used to produce and polymerize hydrogel beads at high-throughput^24^. However, the use of fluorescent dyes to create spectral codes in most cases has limited the number of unique spectral codes to < 100^25,26^. Moreover, fluorescently encoded microspheres are often incompatible with harsh organic solvents, precluding their use in a variety of solid-phase synthesis applications.

Our group recently developed a novel technology (MRBLEs, for <u>M</u>icrospheres with <u>R</u>atiometric <u>B</u>arcode <u>L</u>anthanide <u>E</u>ncoding) that spectrally encodes beads via the ratiometric incorporation of lanthanide nanophosphors (Lns)^27^. Lns have narrow and well-separated emission spectra, making it theoretically possible to generate code sets with 10^5^-10^6^ unique members^28^, and we previously demonstrated the ability to produce and discriminate > 1,100 codes^27^. In addition, MRBLEs can be subsequently functionalized for on-bead solid-phase peptide synthesis^29,30^. However, prior MRBLEs production pipelines were complex, requiring two-layer microfluidic devices with integrated valves and extensive custom pneumatics control hardware^27,28^, limiting widespread adoption. In addition, production throughput was slow (requiring 1.5 hours to produce ∼10,000 beads with a given embedded code), making bead generation for assay development inefficient and labor-intensive.

Here, we present a simple method (MRBLEs 2.0) for efficient high-throughput generation of spectrally encoded and magnetic beads bearing various functional groups for downstream chemical coupling or on-bead synthesis. MRBLEs 2.0 relies on manual mixing of Ln/polymer mixtures followed by droplet generation using single-layer, parallel flow-focusing (FF) devices and off-chip batch polymerization of droplets into beads, thereby simplifying and enhancing throughput of bead production. To facilitate downstream analyte coupling, MRBLEs 2.0 localizes copolymers bearing functional groups typically used for bioconjugation to the surface of the hydrogel matrix during droplet generation and covalently crosslinks these polymers in place during bead polymerization. Using the MRBLEs 2.0 pipeline, we demonstrate the ability to generate monodisperse MRBLEs (CV<7%) containing 48 distinct well-resolved spectral codes (<0.01% probability of code-misassignment) in high-throughput (>∼3,000,000 beads with diameter ∼50 μm within 20 min per code, including washing steps). To highlight the utility of these MRBLEs in downstream assays, we conjugate amine-functionalized oligonucleotides and entire proteins to MRBLEs bearing carboxyl groups with estimated bead loading densities of 10^7^-10^8^ molecules/bead, and additionally conjugate biotin molecules to amine-functionalized MRBLEs, demonstrating the feasibility of direct on-bead peptide synthesis^29,30^. To facilitate bead separation and washing during assays, we additionally show that spectrally encoded beads can also be magnetized via simultaneous incorporation of magnetic nanoparticles with little effect on spectral codes. Finally, for potential assays requiring large numbers of beads, we use droplet splitting to exponentially amplify bead production within the same microfluidic chip footprint. We anticipate that MRBLEs 2.0 will be broadly useful pipeline for a variety of bioassays designed to detect DNA hybridization, protein-protein/peptide interactions, and screen polymers for useful bioactivity.

## Results

### High-throughput production of MRBLEs using a novel parallel flow focuser and syringe pumps

MRBLEs 2.0 requires only 2 syringe pumps, a low-cost microscope, a flood UV illumination source, and an easily fabricated single-layer microfluidic device. To allow for parallel microfluidic droplet generation, the device uses PEEK tubing as ‘jumper cables’ to provide three-dimensional routing of flow channels (see **Figure S1** and Supplemental Methods) without a need for complex multilayer PDMS device fabrication or laser ablation to create connections between layers^31^.

The pipeline produces MRBLEs in 3 stages as shown in **Figure 1a**: (1) off-chip generation of polymer/lanthanide mixtures that will ultimately comprise MRBLEs (‘polymer mixing’), (2) on-chip production of polymer/lanthanide droplets using parallel microfluidic flow focusers (‘droplet production’), and (3) off-chip polymerization of polymer/lanthanide droplets into solid MRBLEs via exposure to UV light (‘bead polymerization’). In the first stage (‘polymer mixing’), ratiometric mixtures of Lns (typically 24, 48, or 96 distinct mixtures each corresponding to a unique spectral code) are combined with a polymer solution off-chip via manual or robotic pipetting and deposited into a standard multi-well plate. At this point, polymer solutions can be stored at 4°C in the dark for weeks prior to MRBLEs synthesis. Immediately before MRBLE droplet production, a UV-activated photoinitiator (lithium phenyl-2,4,6-trimethylbenzoylphosphinate (LAP) is added to each encoded polymer solution to facilitate subsequent off-chip bead polymerization.

**Figure 1.**
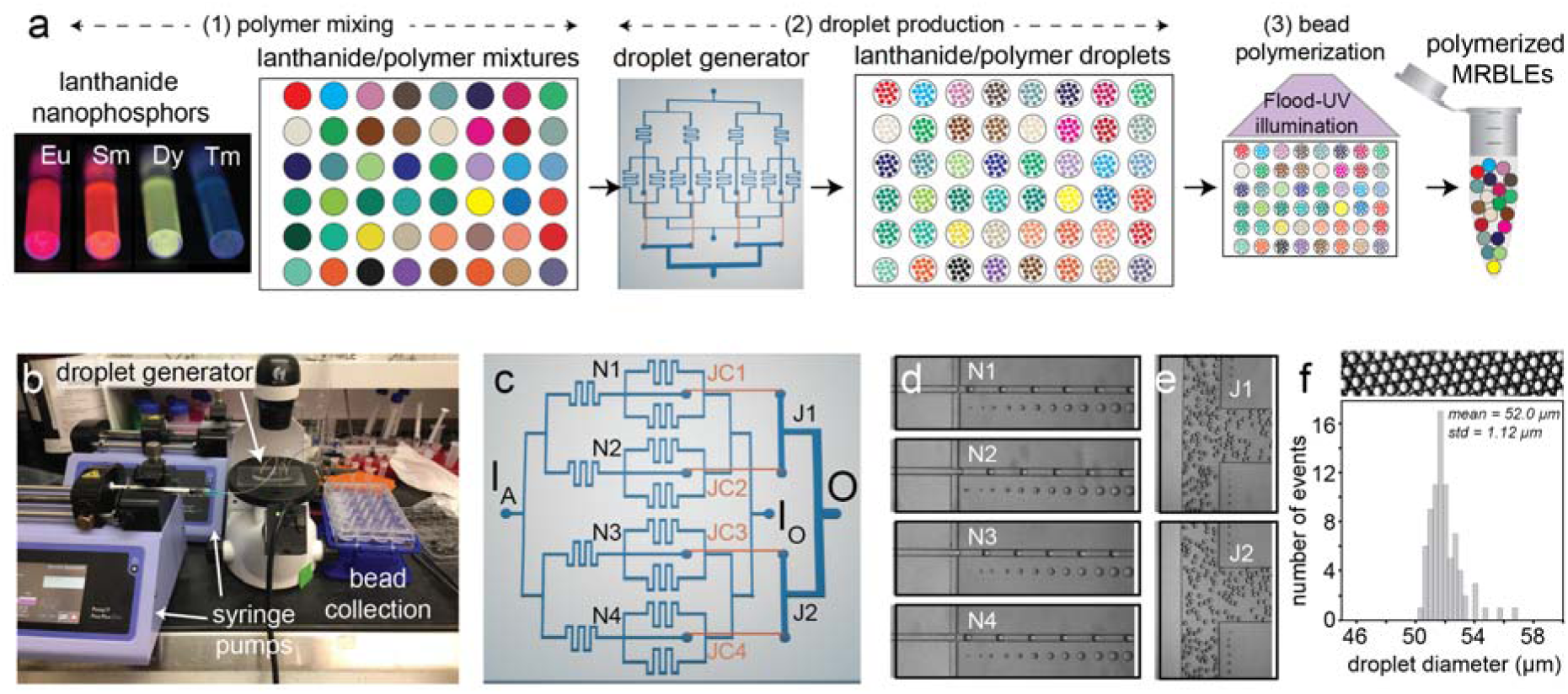
MRBLEs 2.0 high-throughput bead production. **(a)** Overall experimental pipeline: (1) ‘polymer mixing’: ratiometric combinations of lanthanide nanophosphors (each of which comprises a unique spectral code) are mixed with aqueous polymer in individual wells of a multi-well plate; (2) droplet production: mixtures are iteratively introduced into a single-layer microfluidic droplet generation device; (3) bead polymerization: droplets corresponding to each code are collected after synthesis and are exposed to UV light to drive polymerization into solid beads. **(b)** Low-cost setup for high-throughput MRBLEs synthesis consisting of 2 syringe pumps and an inexpensive inverted microscope. **(c)** Schematic of single layer microfluidic device. Aqueous polymer and oil solutions are introduced at I_A_ and I_O_ and meet at 4 flow-focusing nozzles (N1-N4); produced droplets are routed from 4 junctions to 2 additional junctions (J1-2) via ‘jumper cable’ tubing (JC1-4) prior to collection from the droplet outlet (O). **(d)** Images of droplet generation at the 4 flow-focusing nozzles (N1-4). **(e)** Images of droplets merging at the junctions J1 and J2 after passing through ‘jumper cables’. **(f)** Representative image (top) and histogram (bottom) showing measured diameters for produced droplets (diameter = 52.0 ± 1.12 µm, CV=2%, 79 droplets).

In the second stage (‘droplet production’), these Ln mixtures and a fluorinated oil solution (HFE7500 + 2% w/w Ionic Krytox surfactant) are simultaneously introduced into a microfluidic droplet generator using syringe pumps at specific volumetric flow rates (**Figure 1b)**. Within the device, these aqueous polymer and fluorinated oil streams meet at 4 parallel droplet generation nozzles, where changes in interfacial tension cause the aqueous polymer to break off and form monodisperse droplets within the oil stream (**Figures 1c&d**; **Movie S1**). ‘Jumper cables’ route droplets produced at each nozzle to 2 junctions (**Figures 1c&e**) and subsequently to a single output, enhancing production rates 4-fold while allowing collection of all droplets containing a given code within a single well of the 24-well plate. Droplet sizes can be tubed by simply varying the flow rates of aqueous and oil phases (**Figure S2**). Here, we used flow rates of 600 µL/hour (aqueous) and 3200 µL/hour (oil) to generate stable droplets with a measured median diameter of ∼52 µm and an overall CV of 2% (**Figure 1f**). Between codes, the entire device can be flushed with oil and the aqueous tubing (**Figure S1b**, assembled in step 1) can be flushed with water to prevent cross-contamination.

In the final stage (‘bead polymerization’), all droplets from a given code are transferred to a single well of a multiwell plate and the entire plate is exposed to flood UV illumination, allowing simultaneous polymerization of all droplets into solid beads. This pipeline typically produces 3,000,000 MRBLEs/20 min, enhancing throughput by >1,000-fold compared to prior production methods^27^. Polymerized MRBLEs remain highly monodisperse, with a measured median diameter of 52 µm and an overall CV of 7% (**Figure 2a and Figure S3**). This CV is slightly higher than that seen with prior lower-throughput production methods (7% vs 5%, respectively), likely due to a combination of oxygen inhibition and overheating that can induce droplet breakage during off-chip polymerization. However, the benefits associated with enhanced throughput likely outweigh the deleterious effects of a small increase in CV for all but the most sensitive applications.

**Figure 2.**
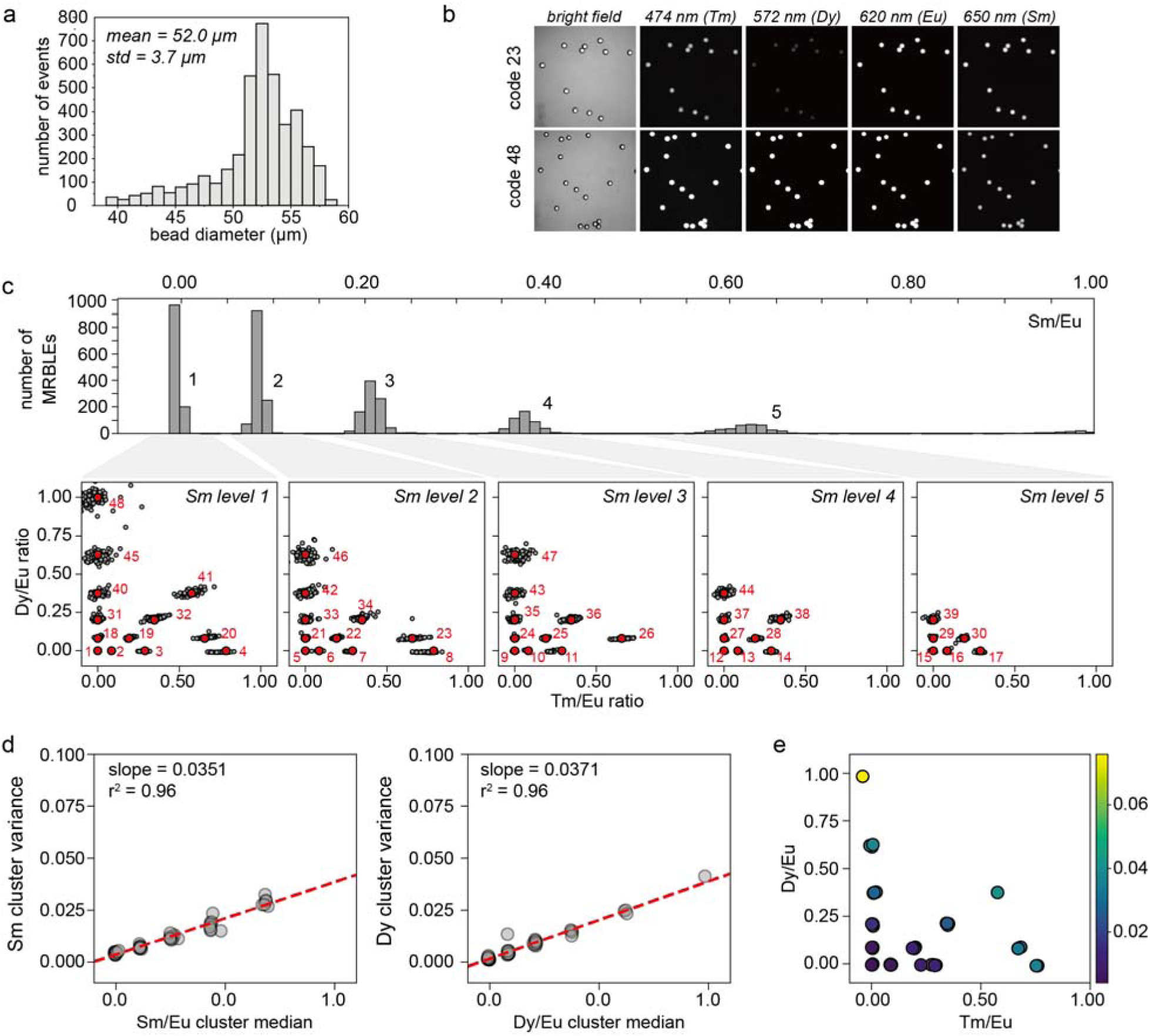
MRBLEs 2.0 spectral codes and code cluster characterization. **(a)** Histogram showing measured diameters for polymerized MRBLEs (diameter = 52.0 + 3.7 µm, CV=7%, 4138 beads). **(b)** Representative bright field and lanthanide emission images of MRBLEs 2.0 beads for 2 different example spectral codes (23 and 48). **(c)** Histogram of Sm bead intensities (top) and scatter plots showing measured Dy/Eu and Tm/Eu ratios for each Sm intensity level (bottom) for 48 code clusters (4138 beads in total). **(d**) Sm/Eu (left) and Dy/Eu (right) cluster variance as a function of cluster median. **(e)** Tm cluster variance as a function of Dy/Eu and Tm/Eu cluster median.

### Spectral codes within MRBLEs produced in high-throughput remain easily resolved from one another

Using MRBLEs in downstream multiplexed assays requires the ability to distinguish embedded codes from one another with high confidence. To test the robustness of spectral encoding and determine the maximum likely coding capacity achievable with high-throughput production, we designed a target matrix of 48 distinct codes comprised of ratios of Dysprosium (Dy), Samarium (Sm), Thulium (Tm), and Europium (Eu) Lns (Dy/Eu, Sm/Eu, and Tm/Eu). We then mixed these ratios off-chip via simple manual pipetting, generated droplets from these mixtures, polymerized the droplets, washed extensively with solvents (to remove unpolymerized material), and imaged the resultant MRBLEs to quantify observed Ln intensity ratios and compare these measured ratios to the desired target ratios.

Images of MRBLEs excited in the deep UV (292 nm) with emission collected across 9 Ln emission channels (435, 474, 536, 546, 572, 620, 630, 650, and 780 nm) established that beads composed of a 21.4% v/v poly(ethylene glycol) diacrylate (PEG-DA) matrix were homogeneously polymerized after UV flood exposure without detectable Ln aggregation (**Figure 2b**). However, we also observed that higher concentrations of PEG-DA led to dramatic aggregation of Lns (**Figure S4**)^27^, likely due to PEG-DA absorption of water molecules required to maintain charge-charge repulsion of polyacrylic acid-wrapped Lns.

In prior work, we observed that emission spectra for Sm are largely orthogonal to those of other Lns such that observed Sm/Eu intensities depend only on the amount of Sm incorporated within each bead^27,28^. Consistent with this, we observed 5 clearly separable Sm/Eu ratios that correspond directly to the intended Sm/Eu target levels (**Figure 2c**, top). By considering only beads at a given Sm/Eu ratio and plotting the observed Dy/Eu ratio against the observed Tm/Eu ratio for each bead, we can further directly visualize individual code clusters and compare each cluster to its intended target value (**Figure 2c**, bottom). All 48 code clusters are present and easily distinguished from one another with the median observed cluster intensity ratio well-centered on the desired target. For Sm/Eu and Dy/Eu ratios, the observed cluster variance depended linearly on the median Sm/Eu or Dy/Eu ratio, respectively (**Figure 2d**), with a slope of ∼0.04 for both Lns. By contrast, the variance for clusters in the Tm/Eu channel depends on both the median Tm/Eu and Dy/Eu ratios (**Figure 2e**), reflecting the fact that Dy and Tm both emit light in the 474 nm emission channel, contributing to crosstalk (**Figures S5a&b**). This measured cluster variance as a function of median value suggests that Sm, Dy, Tm, and Eu could be used to produce well-resolved code sets with 208 clusters separated by 5 standard deviations between each cluster or even larger code sets with smaller separations (839 or 380 clusters at 3 and 4 standard deviation separations, respectively) (**Figure S5c**).

### Direct MRBLEs functionalization during off-chip polymerization

Multiplexed bioassays designed to screen for binding interactions require the ability to couple ‘bait’ analytes of interest (*e.g*. DNA oligonucleotides, whole proteins, or chemically synthesized peptides) to the MRBLEs surface at high density and without crosstalk or analyte dissociation over long storage timescales. Covalent coupling is particularly robust and typically relies on two main functional groups: - COOH (carboxyl groups) or -NH_2_ (amine groups) (**Figure 3a**). Carboxyl groups on the bead are compatible with 1-Ethyl-3-(3-dimethylaminopropyl) carbodiimide (EDC) chemistry, which facilitate subsequent covalent coupling of any molecules bearing a free amine group to beads in aqueous/organic solvents (*e.g*. whole proteins bearing an exposed primary amine or amine-functionalized DNA oligonucleotides). Conversely, amine groups displayed on MRBLEs provide convenient handles for covalent coupling of carboxylated molecules, such as solid-phase synthesis of peptides directly on beads via standard Fmoc coupling chemistry.

**Figure 3.**
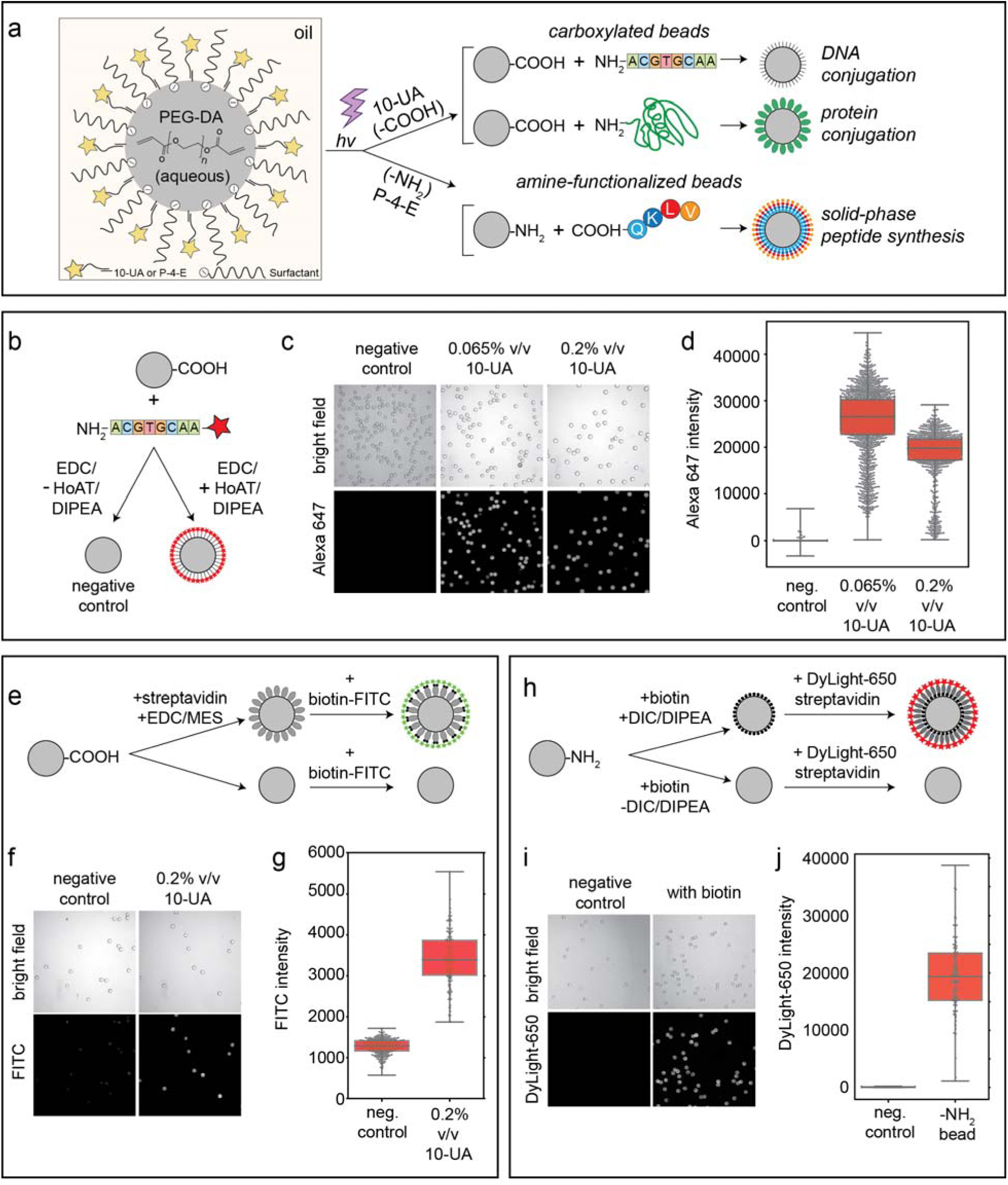
Covalent conjugation of oligonucleotides, proteins, and biotin to MRBLEs co-polymerized with functionalized monomers. **(a)** Schematic illustrating functionalization of PEG-DA hydrogel beads for downstream DNA or protein coupling, or on-bead peptide synthesis. **(b)** Assay used to quantify specific coupling of fluorescently-labeled amine-functionalized oligonucleotides to carboxylated beads. **(c)** Bright field (top) and fluorescence (bottom) images of MRBLEs polymerized in the presence of 0.065% or 0.2% v/v 10-UA comonomers after conjugation with and without required coupling reagents (negative control beads were polymerized with 0.2% v/v 10-UA comonomers). **(d)** Median Alexa-647 intensities for all MRBLEs from each condition. **(e)** Assay used to quantify coupling of streptavidin proteins to carboxylated beads. **(f)** Representative images of MRBLEs incubated with biotin-FITC. **(g)** Median FITC intensities for all MRBLEs from each condition. **(h)** Assay used to quantify on-bead peptide synthesis. **(i)** Representative images of MRBLEs after incubation with DyLight-650-labeled streptavidin. **(j)** Median DyLight-650 intensities for all MRBLEs from each condition.

In prior work, -COOH or -NH_2_ moieties were covalently coupled to the PEG-DA polymer chains comprising the MRBLEs hydrogel matrix via Michael addition after bead synthesis^30^. However, this method was relatively slow (requiring ∼24-48 hours) and susceptible to inhomogeneous distribution of free acrylate groups on bead surface, leading to heterogeneity of coverage during coupling. To address this, we developed a novel functionalization approach that leverages solubility differences between -COOH- or -NH_2_-terminated comonomers and photoinitiator (LAP) to drive polymerization of comonomers at the surface of the hydrogel matrix only (**Figure 3a**). The comonomers used here (10-undecenoic acid (10-UA) for -COOH and pent-4-enylamine (P-4-E) for -NH_2_) are soluble in the HFE 7500 oil, while the LAP is only soluble in water. Exposure of LAP-containing aqueous droplets to UV drives a polymerization chain reaction that begins in the aqueous PEG-DA microsphere core and ultimately reaches the oil/water interface to drive cross-linking of oil-soluble interfacial comonomers to the hydrogel bead matrix (**Figure 3a**). As the concentration of comonomer within the bulk oil phase is very low (0.2% v/v for 10-UA and 0.09% v/v for P-4-E), the chain reaction is then abolished to limit cross-linking to this oil/water interface.

### Direct coupling of oligonucleotides and proteins to MRBLEs and on-MRBLE biotin conjugation

To demonstrate this technique and quantify coupling efficiency and variability for DNA conjugation, we synthesized MRBLEs bearing -COOH functional groups at the surfaces and directly coupled fluorescently labeled, NH_2_-functionalized oligonucleotides to them via EDC chemistry (**Figure 3b**). Images of beads after coupling and washing establish that fluorescence intensities are relatively even across beads (CV ∼27%) and high only when coupling is performed in the presence of HoAT/DIPEA coupling reagents, establishing that coupling is specific (**Figures 3b-d**). Bright field and fluorescence images further establish that beads are undamaged after exposure to coupling reagents and that unbound oligonucleotides are effectively removed by washing (**Figures 3b-d**). Comparison of fluorescence signals for beads polymerized in the presence of 2 different concentrations of 10-UA reveal that intensities are higher but slightly more heterogeneous for 0.065% (v/v) 10-UA, likely reflecting incomplete coverage of the whole droplet surface^32^. Estimation of loading densities by DNA conjugation for the 0.2% (v/v) 10-UA condition suggest ∼10^7^ to 10^8^ oligos bind to the surface of each bead, consistent with prior estimates of MRBLEs peptide loading capacities^30^.

To demonstrate that carboxylated MRBLEs can also be used for covalent coupling of entire proteins under aqueous conditions, we tested the ability to attach streptavidin molecules to MRBLEs surfaces via their primary amines. To visualize attached streptavidin, we then incubated beads with biotin-FITC conjugates, washed 3 times and imaged (**Figure 3e**). As with the DNA conjugation reactions, we observed significant FITC intensities only after covalently coupling streptavidin to surfaces and not in the absence of coupling reagents (**Figures 3f&d**).

Finally, we demonstrated the feasibility of using NH_2_-functionalized MRBLEs for solid-phase peptide synthesis by: (1) polymerizing MRBLEs with P-4-E (0.09%, v/v) added to the oil phase, (2) incubating these functionalized MRBLEs with biotin (which bears a free -COOH group analogous to standard Fmoc-amino acids) in either the presence or absence of required coupling reagents, (3) incubating with DyLight 650-labeled streptavidin, (4) washing, and then (5) imaging to quantify amount of bead-bound fluorescence (**Figure 3h**). As with the other coupling reactions, we observed strong fluorescence only in the presence of coupling reagents, establishing that presented NH_2_ groups can be used for subsequent specific conjugation (**Figures 3i&j**).

### Spectrally encoded magnetic MRBLEs improve separation efficiencies and increase coding capacity

The ability to selectively magnetize particles provides an additional coding axis, thereby increasing the number of codes that can be distinguished from one another (*e.g*. generating the same set of spectral codes in the presence and absence of magnetic particles increases coding capacity 2-fold). In addition, magnetic beads have particular advantages for bead-based separations, improving bead retention during rinsing and removal of excess supernatant via the application of a magnetic field and facilitating more stringent washing. Finally, magnetic beads can also aid in loading beads into microwells for high-throughput experiments (*e.g*. single-cell phenotyping).

To test the ability to create magnetic MRBLEs, we synthesized Fe_3_O_4_ nanoparticles by modifying a previously published co-precipitation method^33^ to include extra PAA in the precursor solution, thereby enhancing control of nucleation and nanoparticle wrapping as well as preventing nanoparticle aggregation. The resultant Fe_3_O_4_ nanoparticles were ∼46 nm in diameter and appeared optically transparent (**Figure S6**). When these Fe_3_O_4_ nanoparticles were incorporated within MRBLEs, all MRBLEs in an Eppendorf tube could be attracted to one side of the tube by simply holding a magnet at the side of the tubing (**Figure 4a** and **Movie S2**).

**Figure 4.**
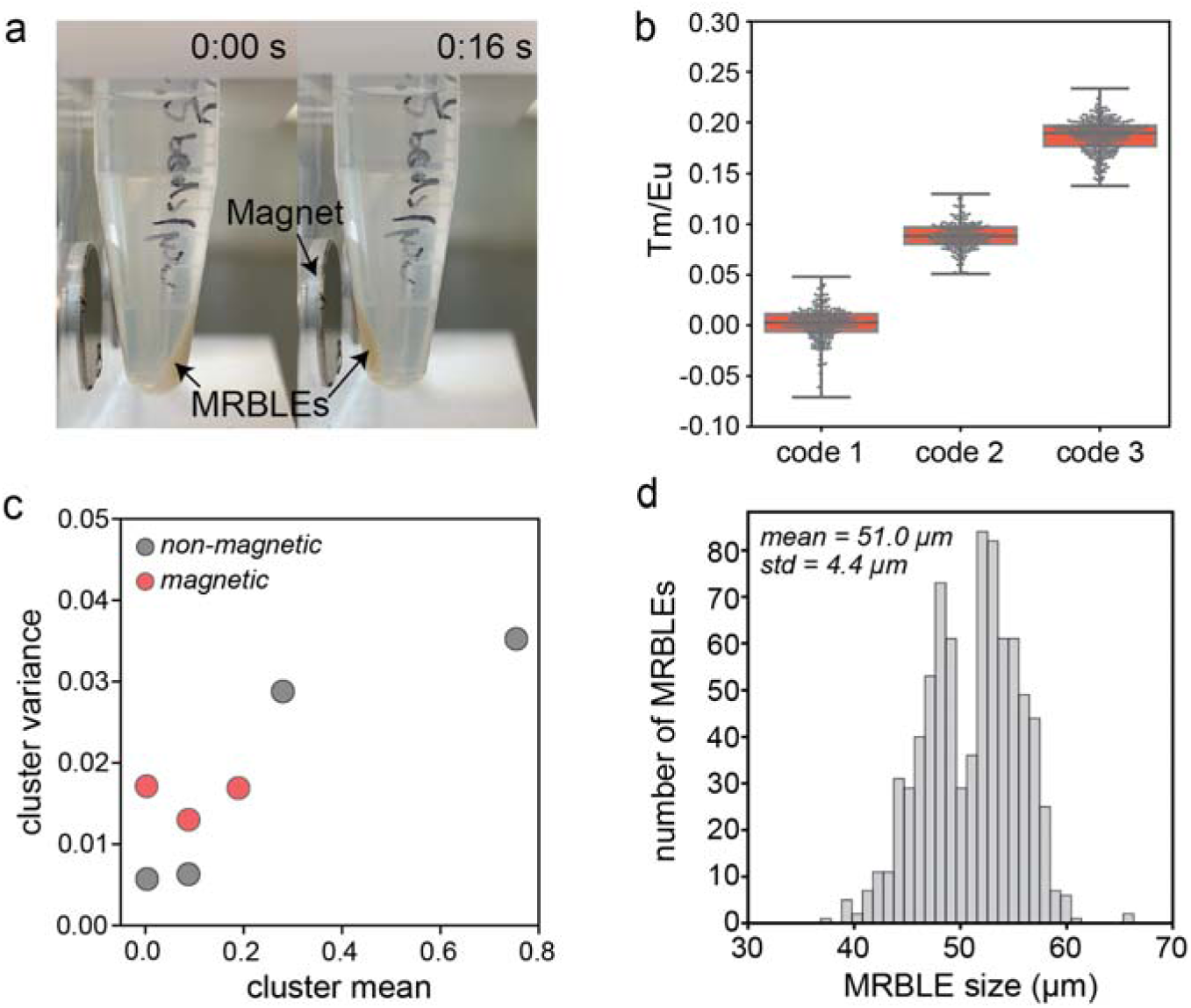
MRBLEs 2.0 beads containing embedded spectral and magnetic codes. **(a)** Images showing magnet-induced movement of MRBLEs 2.0 beads containing magnetic nanoparticles. **(b)** Measured Tm/Eu intensities for MRBLEs containing magnetic nanoparticles and 3 different closely-spaced Tm/Eu ratios. **(c)** Measured cluster variance for Tm/Eu code clusters in the presence and absence of magnetic nanoparticles. **(d)** Measured size distribution for spectrally encoded magnetic MRBLES.

To test if MRBLEs spectral codes can be resolved accurately even in the presence of 10% Fe_3_O_4_ nanoparticles, we synthesized a small library of MRBLEs including a single Ln species. In prior encoding efforts, Tm has consistently been the least efficient Lns emitter^34^ due to relatively low energy transfer efficiency between doped Tm^3+^ and VO_4_^3-^, resulting in weak blue emission^35^. To provide a particularly stringent test of potential coding capacity, we therefore synthesized magnetic MRBLEs containing the 3 lowest Tm/Eu levels used previously (see **Figure 2**), mixed them together, and attempted to resolve the 3 populations (**Figure 4b**). All 3 levels were clearly distinguishable from one another, with a < 0.01% probability of miscalling a code. Comparison of Tm/Eu cluster variance as a function of mean Tm/Eu level further establishes that variances are only ∼2-fold higher in the presence of magnetic nanoparticles (**Figure 4c**), suggesting that up to 1,000-plex spectral code spaces remain possible. As with prior tests, magnetic MRBLEs were fairly monodisperse, with diameters of 51 ± 4 µm (**Figure 4d**, mean ± standard deviation; CV = 7.8%).

### Exponential droplet splitting dramatically increases MRBLE production rates

Leveraging MRBLEs for broad use requires the ability to rapidly and economically produce large numbers of beads. While increasing fluid flow rates can boost droplet production within a narrow range, large increases in flow rates drive a transition in droplet formation between dripping to turbulent jetting regimes, yielding very polydisperse beads. As an alternate approach, throughput can be increased by simply adding additional parallel flow focusers (FFs) to the ‘jumper cable’ synthesis device described above (**Figure 5a**); however, this approach increases device size linearly with throughput (production rates scales as *N*, where *N* represents the number of FFs). To address this, we designed and fabricated an additional device that combined exponential droplet splitting with ‘jumper cables’ to boost droplet production rates without increasing device area. In this device, a large droplet (∼160 μm in lateral diameter and 50 μm in height) is initially split to form two smaller ones when encountering two bifurcated channels (**Figure 5a**). These smaller droplets are subsequently split by more Y-junction structures, thus generating an exponential amplification of bead production (with rates increased by 2^*N*^, where *N* represents the number of splitting cycles) (**Figures 5b&d** and **Movie S3**). As with the prior method, produced droplets are routed to a single outlet via ‘jumper cables’, collected, and polymerized *en masse* via flood UV. For a device with 8 × 4 splitters (8 splitters in each pathway, 4 pathways in total), measured droplet production rates averaged 10^6^/min at aqueous and oil flow rates of 1500 µL/h and 5400 µL/h, respectively, representing a ∼3-fold increase over a linear production device with 4 FFs (**Figure 1b**). Resultant droplets were slightly more polydisperse (CV = 12%) but spectral codes remained easily distinguished from one another (**Figure 5c)**.

**Figure 5.**
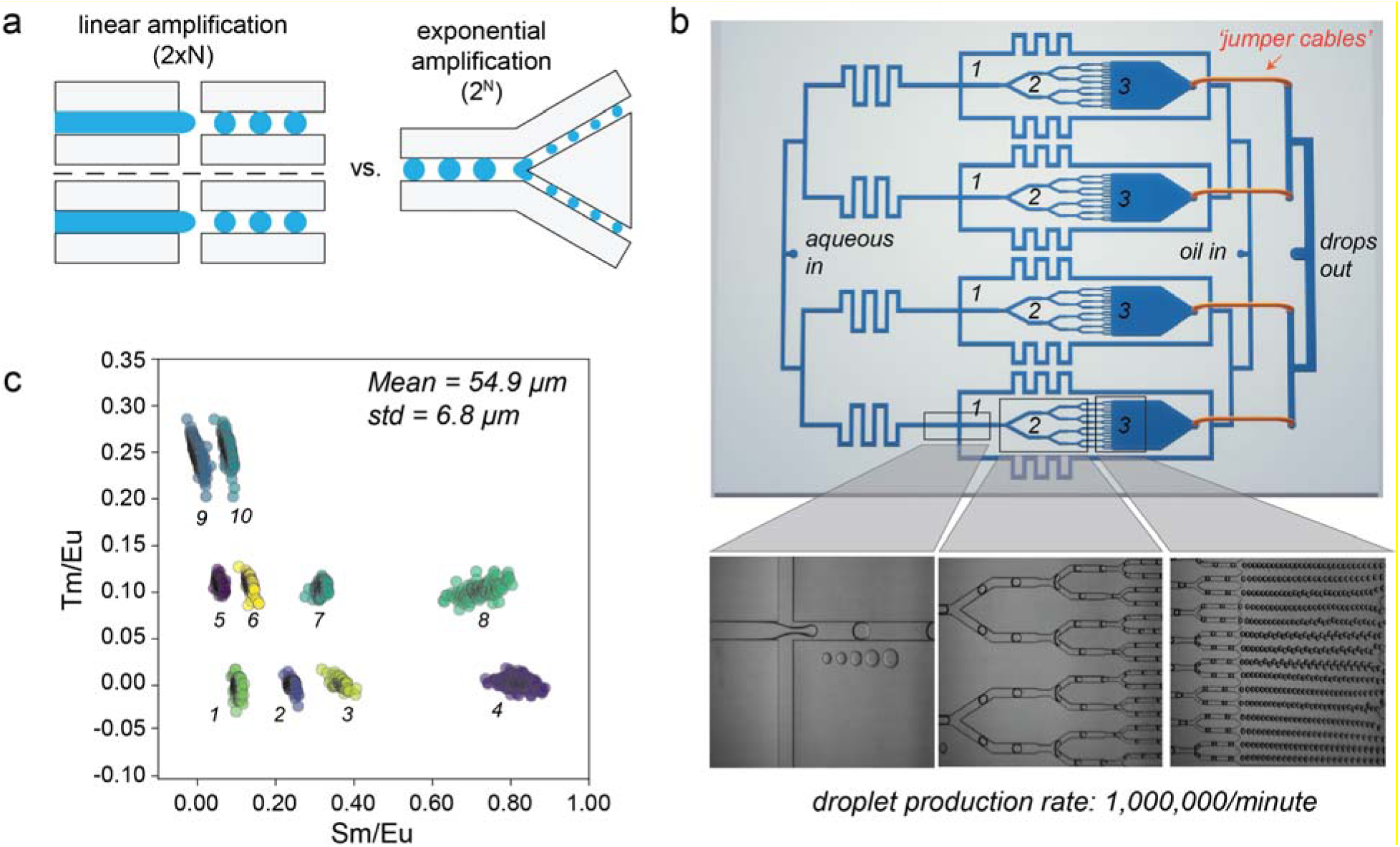
High-throughput MRBLEs 2.0 bead production via exponential droplet splitting. **(a)** Schematic demonstrating linear vs exponential droplet amplification. **(b)** Schematic of device for exponential droplet production including 4 flow focusing nozzles for generating large droplets (*1*) and a series of 4 channel bifurcations (*2*) leading to a large channel for droplet collection (*3*) (top); representative images from these 3 device regions showing droplet formation, splitting, and collection (bottom). **(c)** Measured Tm/Eu and Sm/Eu ratios from a 10-code set produced via exponential droplet splitting with bead size indicated at top left.

## Discussion

Developing multiplexed bead-based assays with the potential for broad translation and adoption requires the ability to produce large batches of spectrally encoded beads at low cost. Large bead batches make it possible to distribute beads widely for use in other labs during initial assay development and optimization and further minimize experimental noise resulting from batch-to-batch variation. Increasing the number of droplet generators within a device provides an obvious way to boost production rates, and this approach has been used successfully for a variety of different glass or PDMS droplet generation devices (including co-flow nozzles^36^, T-junctions^37,38^, flow-focusers^39-44^, and discontinuous step generators^37,38^). However, the need to route fluidic channels containing dispersed and continuous phases to and from many different droplet generating nozzles without channels intersecting one another presents fabrication challenges. In the past, this has been addressed by fabricating complex 3D wafer designs (for discontinuous step generators) or assembling devices from multiple PDMS layers with laser-cut or punched ‘vias’ connecting channels between layers (for T-junctions and flow-focusers). Here, we adapt prior work that used tubing to connect different PDMS functional ‘modules’^45,46^ to create ‘jumper cables’ that connect different channels within a single device architecture. This strategy significantly reduces the fabrication complexity required to flexibly route channels in 3 dimensions while still minimizing fluidic dead volumes. Using this strategy, we demonstrated the ability to produce 150,000 MRBLEs/min (linear amplification) or 450,000 MRBLEs/min (exponential amplification), representing a pellet volume of ∼1 mL/hour or 3 mL/hour and a 1000- to 3000-fold increase in throughput from prior work^27^.

The utility of spectrally encoded beads in downstream assays also depends critically on the ease with which analytes can be coupled to beads. Streptavidin or antibody-conjugated beads are often used to recruit and display biotinylated or epitope-tagged proteins, but these interactions have non-zero dissociation rates, leading to the possibility of analyte loss and rebinding to drive cross-contamination during long-term storage. The ability to covalently link molecules of interest to encoded beads bearing functional groups widely used in bioconjugation chemistry sidesteps this issue, but can be difficult to implement with many commonly used fluorescently labeled bead matrices. Here, we demonstrated the ability to incorporate functional groups by simply generating aqueous polymer droplets in an oil stream containing functionalized co-monomers bearing carboxyl or amino groups. This same approach could easily be extended to a variety of other functional groups^47^, including azides for downstream ‘click’ chemistry, hydroxyl groups for cyanogen bromide-activated coupling of proteins, hydrazide groups for oxidized carbohydrate proteins and chloromethyl groups for coupling NH_2_ groups in proteins or other biomolecules. In addition, this approach could facilitate straightforward coupling of multiple molecules of interest to a single bead (*e.g*. proteins and oligonucleotides simultaneously) by adding multiple comonomers to the oil phase during droplet generation, facilitating production of beads for single-cell profiling methods such as CITE-seq^48^ or ECCITE-seq^49^.

The simultaneous magnetic and spectral encoding strategy demonstrated here enhances the ability to remove a portion of molecules after coupling or synthesis to quantitatively assess bead loading or synthesis quality. As an example, peptides could be synthesized directly on a mixture of spectrally encoded beads and unencoded magnetic beads, making it possible to remove magnetic beads after synthesis for peptide cleavage and investigation via mass spectrometry.

In future work, the simple encoded bead production and functionalization pipeline presented here could be integrated with various liquid handling robots and high-throughput droplet generation technologies for a variety of downstream commercial and translational applications. Lns, PEG-DA polymer, functionalized comonomers, and photoinitiators are all commercially available and relatively inexpensive, facilitating development of low-cost assays. In addition, the ability to mix Lns and polymer before introducing solutions into devices for droplet generation makes it possible to automate this process for production of large code sets using commercial liquid handling robots capable of mixing viscous solutions (*e.g*. the LabCyte Echo or Bravo liquid handling systems). These mixed solutions could then be introduced into devices optimized for ultra-high-throughput droplet production via centrifugal methods^50-52^, in-air jetting droplet generators^53^, or glass-silicon chips recently shown to produce droplets at rates exceeding 5.5 billion droplets/minute^39^. In this way, production of lanthanide-encoded MRBLEs at industrial scale could become feasible, opening up a wide range of new bead-based multiplexed assays.

## Materials and Methods

### Device design, photolithography, and fabrication

All devices used in this paper were designed and fabricated via standard soft lithography protocols^54^. Briefly, microfluidic molding masters were created by: (1) coating 4” test-grade silicon wafers (University Wafer, South Boston, MA) with a single layer of SU-8 2050 negative photoresist, (2) soft baking, (3) exposing this SU-8 layer to UV light passing through a printed transparency mask (designed in AutoCAD (Autodesk); printed at 50,000 dpi by Fineline Imaging), (4) post-exposure baking, and (5) developing away uncured photoresist using SU-8 developer (Microchem Corp, Newton, MA) according to standard manufacturer instructions. These molding masters were then used to cast single-layer droplet generators composed of a 1:5 ratio of poly(dimethylsiloxane) crosslinker:base (PDMS, RTV 615, Momentive Performance Materials, Albany, NY), and these resultant devices were assembled as described in Supplementary Methods. All the holes were punched by catheter punch (SYNEO, 0.025” ID × 0.035” OD, Part No: CR0350255N20R4) to fit the outer diameters of PEEK tubing (ZEUS, 0.010” ID × 0.020” OD) and steel blunt pins (New England Small Tube, Part No: NE-1310-02). All design files and detailed protocols for molding master and device fabrication are available as Supplementary Files and in an associated OSF repository (https://osf.io/jvnpc/).

### Bead synthesis

Mixtures of PEG-DA polymer, lanthanides, and LAP photoinitiator were prepared largely as described previously^27^. In general, pre-mixed formulas (250 μL in total) were generated by varying ratios of three monomer master mixtures each containing different Ln nanophosphors. All aqueous master mixtures contained purified water with 21.4% v/v PEG-DA (Sigma-Aldrich, average Mn 700) and 5% v/v YVO_4_:Eu (50 mg/mL). The “Dy”, “Sm” and “Tm” master mixtures also contained 21.3% v/v YVO_4_:Dy (50 mg/mL), 21.3% v/v YVO_4_:Sm (50 mg/mL) and 21.3% v/v YVO_4_:Tm (50 mg/mL), respectively. Each formula was prepared based on the reference ratio listed in **Table S1**. Directly prior to injection of polymer solutions into the device, we added 3% v/v photoinitiator Lithium phenyl-2,4,6-trimethylbenzoylphosphinate (Sigma-Aldrich, “LAP”, 39.2 mg/mL in DI water). Produced Ln/polymer droplets were collected through Tygon tubing into a 24-well plate (Thermo Fisher Scientific). To prevent premature evaporation of HFE7500 and resultant droplet breakage during initial droplet production, we filled each well with ∼80 μL of oil solution prior to droplet generation. To polymerize these Ln/polymer droplets into solid MRBLEs beads, we flood-exposed wells to UV light (IntelliRay, UV0338) for 2 minutes at 100% amplitude (7” away from the lamp, power= ∼50-60 mW/cm^2^). We recommend direct UV polymerization after the production of every 2 codes to reduce any potential droplet breakage. After polymerization, we washed beads with 2 mL dimethylformamide (DMF, Thermo Fisher Scientific) for 20 s, 2 mL dichloromethane (DCM, Thermo Fisher Scientific) for 10 s and 2 mL methanol (Thermo Fisher Scientific) for 20 s, and then resuspended in either 1 mL 1X phosphate buffered saline (PBS) (Thermo Fisher Scientific) with 0.01% (v/v) Tween-20 (Sigma-Aldrich) (PBST) for the aqueous EDC chemistry and biotinylation of NH_2_-MRBLEs or dimethyl sulfoxide (DMSO, Thermo Fisher Scientific) for oligonucleotide conjugation. For making magnetic MRBLEs, we added 10% v/v of the Fe_3_O_4_ nanoparticle solution (150 mg/mL) to each recipe. Detailed protocols for synthesis of Lns and magnetic nanoparticles are provided in SI Methods.

### Oligonucleotide Conjugation

To prepare for oligonucleotide conjugation, beads were rinsed 3 times with 200 µL of DMSO by spinning down the beads into a pellet and removing the supernatant. After the final spin, we resuspended the beads in DMSO at a concentration of 10000 beads per 170 µL volume reaction. We assembled the ‘conjugation reaction’ by combining: 20 µL 1-ethyl-3-(3-dimethylaminopropyl)carbodiimide (EDC) from a 300 mM stock solution prepared by dissolving 58 mg EDC in 1 mL of DMSO; 20 µL 1-hydroxy-7-azabenzotriazole (HOAT) from a 60 mM solution prepared by dissolving 9.2 mg of HOAT in 1 mL of DMSO; and 20 µL diisopropylethylamine (DIPEA) from a 300 mM solution prepared by adding 32 µL DIPEA to 968 µL DMSO. We then incubated beads in this EDC, HOAT, and DIPEA solution for 15 minutes on a shaker (or rotator at 15 rpm) to ensure adequate mixing. After 15 minutes, we added 10 µL of a 100 µM stock solution of oligonucleotides modified at the 5’ end with amines and a carbon spacer (the “5AmMC12” modification from Integrated DNA Technologies) and modified at the 3’ end with an Alexa-647 dye and an additional 20 µL each of the EDC, HOAT, and DIPEA^55^. We then incubated the reaction at room temperature for 16 hours on a shaker. After this incubation, we neutralized the conjugation reaction by adding 50 µL of a 500 mM ethanolamine solution (prepared by adding 30.2 µL ethanolamine to 968 µL DMSO) to each reaction and incubating for 1 hour. Following conjugation, we washed beads three times with 200 µL PBS containing 0.1% (v/v) Tween 20. We then combined all beads into a single Eppendorf, resuspended at 100 beads/µL in PBS with 0.1% (v/v) Tween 20, and stored at 4 °C.

### Aqueous EDC chemistry

For streptavidin-coated MRBLEs, we washed 150 μL of carboxy MRBLEs and resuspended them in 200 μL of MES buffer (pH = 4.5) supplemented with 0.01% (v/v) Tween 20. Next, we added 200 μL of a freshly made 1-Ethyl-3-(3-dimethylaminopropyl)carbodiimide (EDC, Sigma-Aldrich) solution 2% w/v (corresponds to 10 mg EDC in 500 μL MES buffer) into the bead solution and incubated the entire reaction for 3.5 hours at room temperature on a rotator. The O-acylisourea intermediate on the beads is unstable in aqueous solutions, thus the bead mixture was subsequently immediately resuspended in 400 μL of borate buffer (pH = 8.5) supplemented with 0.01% (v/v) Tween 20. To conjugate streptavidin, we added 16 μL of 1 mg/mL streptavidin (Sigma-Aldrich) in borate buffer into the mixture and rotated the whole slurry overnight at 4°C. After incubation, we quenched the reaction by adding 10 μL of 0.25 M ethanolamine in borate buffer and incubating on a rotator for 30 minutes at 4°C. The final product was washed 3 times and resuspended in 200 μL PBST buffer for further use. To test the conjugation efficiency, we incubated 20 μL streptavidin MRBLEs with 0.5 μL FITC-biotin (Thermo Fisher Scientific, 10 mg/mL in DMSO) for 1h on an end over end rotator at 30 rpm, washed with 1 mL PBST 3 times, and then imaged. In our experience, beads can be stored at 4 °C for ∼ 6 months without a loss of streptavidin binding efficiency.

### Biotinylation of NH_2_-MRBLEs

MRBLEs coated with amine groups using 0.09% (v/v) pent-4-enylamine were stored in PBST and extensively washed with DCM, methanol, and DMF prior to biotin conjugation. For each conjugation reaction, we combined ∼10,000 beads with 39 mg Biotin, 24 µl N,N’-Diisopropylcarbodiimide (DIC) and 56 µl DIPEA in 400 μL DMF twice overnight, rotating at room temperature. After conjugation, we washed beads serially with 1 mL each o fDMF, methanol, DCM, DMF, water and PBST. After washing, we passivated ∼2,000 beads with 5 % BSA PBST for one hour at room temperature on a rotator at 30 rpm; the other ∼8000 beads were stored at 4°C. We then exchanged the 5 % BSA PBST solution to 2 % BSA in PBST via washing and resuspension, added 1 µL of 1 mg/mL DyLight 650 tagged streptavidin (Abcam), and incubated for 30 minutes at 4°C on a rotator at 30 rpm. Finally, we washed all beads 3 times with PBST and imaged as described below.

### Bead imaging and data analysis

Bead imaging was performed on a Nikon Ti microscope with a custom UV transilluminator largely as described previously^28^. Briefly, we imaged lanthanide emission at 9 distinct wavelengths using 9 emission filters (435/40, 474/10, 536/40, 546/6, 572/15, 620/14, 630/92, 650/13, and 780/20 nm) and captured images using an sCMOS camera (Andor) (Andor Technology plc., Belfast, Northern Ireland). We then extracted the most probable Ln ratios associated with each bead via linear unmixing relative to a series of Ln reference spectra^34^. Binding of Alexa-647 DNA oligonucleotides, FITC-biotin, and DyLight-650 streptavidin was visualized using Cy5, eGFP, and Cy5 filter cube sets, respectively.

## Acknowledgements

This work was supported by NIH grants 1DP2GM123641 and R01GM107132. P. M. F. is a Chan Zuckerberg Biohub Investigator and acknowledges the support of a Sloan Research Foundation Fellowship. Y.F. is a Cancer Research Institute Postdoctoral Fellow supported by the Cancer Research Institute Postdoc Fellowship. A.K.W was funded by Natural Sciences and Engineering Research Council of Canada Postdoctoral Fellowship. J.B.H. was funded by grant NNF17OC0025404 from the Novo Nordisk Foundation and the Stanford Bio-X Program. Part of this work was performed at the Stanford Nano Shared Facilities (SNSF), supported by the National Science Foundation under award ECCS-1542152. The authors would also like to thank D. Chan for helpful discussions concerning polymers and K. Brower for guidance with spectrally encoded bead synthesis.

## Author contributions

Y.F. and P.M.F. conceptualized the platform and validation experiments; Y.F. designed and built the ‘jumper cable’ platforms, synthesized magnetic nanoparticles and performed the aqueous EDC chemistry and streptavidin-coated MRBLEs imaging; Y.F., A.K.W. and J.B.H generated MRBLEs; A.K.W. imaged 48 code-sets and performed oligonucleotide conjugation experiments; J.B.H. conjugated biotin to NH_2_-MRBLEs; Y.F., A.K.W., and J.B.H. analyzed data; E.A.A. conceptualized the direct MRBLEs functionalization; and P.M.F. provided funding, resources, mentorship, and project supervision. Y.F., A.K.W., J.B.H., and P.M.F. wrote the paper.

## References

1 Kingsmore, S. F. Multiplexed protein measurement: technologies and applications of protein and antibody arrays. Nature reviews Drug discovery 5, 310–321 (2006).

2 Heller, M. J. DNA microarray technology: devices, systems, and applications. Annual review of biomedical engineering 4, 129–153 (2002).

3 Braeckmans, K., De Smedt, S. C., Leblans, M., Pauwels, R. & Demeester, J. Encoding microcarriers: present and future technologies. Nature reviews Drug discovery 1, 447–456 (2002).

4 Finkel, N. H., Lou, X., Wang, C. & He, L. Peer Reviewed: Barcoding the Microworld. Analytical Chemistry 76, 352 A–359 A, doi:10.1021/ac0416463 (2004).

5 Wilson, R., Cossins, A. R. & Spiller, D. G. Encoded microcarriers for high-throughput multiplexed detection. Angewandte Chemie International Edition 45, 6104–6117 (2006).

6 Birtwell, S. & Morgan, H. Microparticle encoding technologies for high-throughput multiplexed suspension assays. Integrative Biology 1, 345–362 (2009).

7 Broder, G. R. et al. Diffractive micro bar codes for encoding of biomolecules in multiplexed assays. Analytical chemistry 80, 1902–1909 (2008).

8 Cederquist, K. B., Dean, S. L. & Keating, C. D. Encoded anisotropic particles for multiplexed bioanalysis. Wiley Interdisciplinary Reviews: Nanomedicine and Nanobiotechnology 2, 578–600 (2010).

9 Lawrie, G. A., Battersby, B. J. & Trau, M. Synthesis of optically complex core–shell colloidal suspensions: pathways to multiplexed biological screening. Advanced Functional Materials 13, 887–896 (2003).

10 Lee, H., Kim, J., Kim, H., Kim, J. & Kwon, S. Colour-barcoded magnetic microparticles for multiplexed bioassays. Nature materials 9, 745 (2010).

11 Kim, L. N. et al. Shape-encoded silica microparticles for multiplexed bioassays. Chemical Communications 51, 12130–12133 (2015).

12 Chapin, S. C., Appleyard, D. C., Pregibon, D. C. & Doyle, P. S. Rapid microRNA profiling on encoded gel microparticles. Angewandte Chemie International Edition 50, 2289–2293 (2011).

13 Appleyard, D. C., Chapin, S. C. & Doyle, P. S. Multiplexed protein quantification with barcoded hydrogel microparticles. Analytical chemistry 83, 193–199 (2010).

14 Zhang, F. et al. Fluorescence Upconversion Microbarcodes for Multiplexed Biological Detection: Nucleic Acid Encoding. Advanced Materials 23, 3775–3779, doi:10.1002/adma.201101868 (2011).

15 Zhang, F. et al. Rare-Earth Upconverting Nanobarcodes for Multiplexed Biological Detection. Small 7, 1972–1976, doi:10.1002/smll.201100629 (2011).

16 Purohit, S. et al. Multiplex glycan bead array for high throughput and high content analyses of glycan binding proteins. Nature Communications 9, 258, doi:10.1038/s41467-017-02747-y (2018).

17 Fulton, R. J., McDade, R. L., Smith, P. L., Kienker, L. J. & Kettman, J. R. Advanced multiplexed analysis with the FlowMetrixTM system. Clinical chemistry 43, 1749–1756 (1997).

18 Houser, B. Bio-Rad’s Bio-Plex® suspension array system, xMAP technology overview. Archives of Physiology and Biochemistry 118, 192–196, doi:10.3109/13813455.2012.705301 (2012).

19 Waterboer, T., Sehr, P. & Pawlita, M. Suppression of non-specific binding in serological Luminex assays. Journal of Immunological Methods 309, 200–204, doi: https://doi.org/10.1016/j.jim.2005.11.008 (2006).

20 Le Goff, G. C., Srinivas, R. L., Hill, W. A. & Doyle, P. S. Hydrogel microparticles for biosensing. European polymer journal 72, 386–412, doi:10.1016/j.eurpolymj.2015.02.022 (2015).

21 Ullah, F., Othman, M. B. H., Javed, F., Ahmad, Z. & Akil, H. M. Classification, processing and application of hydrogels: A review. Materials Science and Engineering: C 57, 414–433, doi: https://doi.org/10.1016/j.msec.2015.07.053 (2015).

22 Moorthy, J., Burgess, R., Yethiraj, A. & Beebe, D. Microfluidic Based Platform for Characterization of Protein Interactions in Hydrogel Nanoenvironments. Analytical chemistry 79, 5322–5327, doi:10.1021/ac070226l (2007).

23 Nallur, G. et al. Protein and Nucleic Acid Detection by Rolling Circle Amplification on Gel-based Microarrays. Biomedical Microdevices 5, 115–123, doi:10.1023/a:1024535110995 (2003).

24 Roh, Y. H., Lee, H. J. & Bong, K. W. Microfluidic Fabrication of Encoded Hydrogel Microparticles for Application in Multiplex Immunoassay. BioChip Journal 13, 64–81 (2019).

25 Dagher, M., Kleinman, M., Ng, A. & Juncker, D. Ensemble multicolour FRET model enables barcoding at extreme FRET levels. Nature Nanotechnology 13, 925–932, doi:10.1038/s41565-018-0205-0 (2018).

26 Liu, J., Sun, L., Zhan, H. & Fan, L.-J. Preparation of Fluorescence-Encoded Microspheres Based on Hydrophobic Conjugated Polymer–Dye Combination and the Immunoassay. ACS Applied Bio Materials 2, 3009–3018, doi:10.1021/acsabm.9b00337 (2019).

27 Nguyen, H. Q. et al. Programmable Microfluidic Synthesis of Over One Thousand Uniquely Identifiable Spectral Codes. Advanced Optical Materials 5, 1600548, doi:10.1002/adom.201600548 (2017).

28 Gerver, R. E. et al. Programmable microfluidic synthesis of spectrally encoded microspheres. Lab on a Chip 12, 4716–4723, doi:10.1039/c2lc40699c (2012).

29 Nguyen, H. et al. Peptide library synthesis on spectrally encoded beads for multiplexed protein/peptide bioassays. Vol. 10061 PWB (SPIE, 2017).

30 Nguyen, H. Q. et al. Quantitative mapping of protein-peptide affinity landscapes using spectrally encoded beads. eLife 8, e40499, doi:10.7554/eLife.40499 (2019).

31 Huft, J., Da Costa, D. J., Walker, D. & Hansen, C. L. Three-dimensional large-scale microfluidic integration by laser ablation of interlayer connections. Lab on a Chip 10, 2358–2365, doi:10.1039/c004051g (2010).

32 Riechers, B. et al. Surfactant adsorption kinetics in microfluidics. Proceedings of the National Academy of Sciences, 201604307, doi:10.1073/pnas.1604307113 (2016).

33 Upadhyay, S., Parekh, K. & Pandey, B. Influence of crystallite size on the magnetic properties of Fe3O4 nanoparticles. Journal of Alloys and Compounds 678, 478–485, doi: https://doi.org/10.1016/j.jallcom.2016.03.279 (2016).

34 Harink, B., Nguyen, H., Thorn, K. & Fordyce, P. An open-source software analysis package for Microspheres with Ratiometric Barcode Lanthanide Encoding (MRBLEs). PLOS ONE 14, e0203725, doi:10.1371/journal.pone.0203725 (2019).

35 An, Z., Xiao, X., Yu, J., Mao, D. & Lu, G. Controlled synthesis and luminescent properties of assembled spherical YPxV1-xO4:Ln3+ (Ln = Eu, Sm, Dy or Tm) phosphors with high quantum efficiency. RSC Advances 5, 52533–52542, doi:10.1039/c5ra08993j (2015).

36 Nisisako, T., Ando, T. & Hatsuzawa, T. High-volume production of single and compound emulsions in a microfluidic parallelization arrangement coupled with coaxial annular world-to-chip interfaces. Lab on a Chip 12, 3426–3435, doi:10.1039/c2lc40245a (2012).

37 Femmer, T. et al. High-Throughput Generation of Emulsions and Microgels in Parallelized Microfluidic Drop-Makers Prepared by Rapid Prototyping. ACS Applied Materials & Interfaces 7, 12635–12638, doi:10.1021/acsami.5b03969 (2015).

38 Sugiura, S., Nakajima, M., Tong, J., Nabetani, H. & Seki, M. Preparation of Monodispersed Solid Lipid Microspheres Using a Microchannel Emulsification Technique. Journal of Colloid and Interface Science 227, 95–103, doi: https://doi.org/10.1006/jcis.2000.6843 (2000).

39 Yadavali, S., Jeong, H.-H., Lee, D. & Issadore, D. Silicon and glass very large scale microfluidic droplet integration for terascale generation of polymer microparticles. Nature Communications 9, 1222, doi:10.1038/s41467-018-03515-2 (2018).

40 Conchouso, D., Castro, D., Khan, S. A. & Foulds, I. G. Three-dimensional parallelization of microfluidic droplet generators for a litre per hour volume production of single emulsions. Lab on a Chip 14, 3011–3020, doi:10.1039/c4lc00379a (2014).

41 Dangla, R., Kayi, S. C. & Baroud, C. N. Droplet microfluidics driven by gradients of confinement. Proceedings of the National Academy of Sciences 110, 853, doi:10.1073/pnas.1209186110 (2013).

42 Romanowsky, M. B., Abate, A. R., Rotem, A., Holtze, C. & Weitz, D. A. High throughput production of single core double emulsions in a parallelized microfluidic device. Lab on a Chip 12, 802–807, doi:10.1039/c2lc21033a (2012).

43 Mulligan, M. K. & Rothstein, J. P. Scale-up and control of droplet production in coupled microfluidic flow-focusing geometries. Microfluidics and Nanofluidics 13, 65–73, doi:10.1007/s10404-012-0941-7 (2012).

44 Li, W., Greener, J., Voicu, D. & Kumacheva, E. Multiple modular microfluidic (M3) reactors for the synthesis of polymer particles. Lab on a Chip 9, 2715–2721, doi:10.1039/b906626h (2009).

45 Lee, S.-A. et al. Spheroid-based three-dimensional liver-on-a-chip to investigate hepatocyte– hepatic stellate cell interactions and flow effects. Lab on a Chip 13, 3529–3537, doi:10.1039/c3lc50197c (2013).

46 Utharala, R., Tseng, Q., Furlong, E. E. M. & Merten, C. A. A Versatile, Low-Cost, Multiway Microfluidic Sorter for Droplets, Cells, and Embryos. Analytical chemistry 90, 5982–5988, doi:10.1021/acs.analchem.7b04689 (2018).

47 Hermanson, G. T. in Bioconjugate Techniques (Third Edition) (ed Greg T. Hermanson) 229–258 (Academic Press, 2013).

48 Stoeckius, M. et al. Simultaneous epitope and transcriptome measurement in single cells. Nature Methods 14, 865–868, doi:10.1038/nmeth.4380 (2017).

49 Mimitou, E. P. et al. Multiplexed detection of proteins, transcriptomes, clonotypes and CRISPR perturbations in single cells. Nature Methods 16, 409–412, doi:10.1038/s41592-019-0392-0 (2019).

50 Ahmed, N., Sukovich, D. & Abate, A. R. Operation of Droplet-Microfluidic Devices with a Lab Centrifuge. Micromachines 7, 161, doi:10.3390/mi7090161 (2016).

51 Chen, Z. et al. Centrifugal micro-channel array droplet generation for highly parallel digital PCR. Lab on a Chip 17, 235–240, doi:10.1039/c6lc01305h (2017).

52 Shin, D.-C., Morimoto, Y., Sawayama, J., Miura, S. & Takeuchi, S. Centrifuge-based step emulsification device for simple and fast generation of monodisperse picoliter droplets. Sensors and Actuators B: Chemical 301, 127164, doi: https://doi.org/10.1016/j.snb.2019.127164 (2019).

53 Visser, C. W., Kamperman, T., Karbaat, L. P., Lohse, D. & Karperien, M. In-air microfluidics enables rapid fabrication of emulsions, suspensions, and 3D modular (bio)materials. Science Advances 4, eaao1175, doi:10.1126/sciadv.aao1175 (2018).

54 Xia, Y. & Whitesides, G. M. Soft Lithography. Angewandte Chemie International Edition 37, 550–575, doi:10.1002/(sici)1521-3773(19980316)37:5<550::aid-anie550>3.0.co;2-g (1998).

55 Li, Y. et al. Optimized Reaction Conditions for Amide Bond Formation in DNA-Encoded Combinatorial Libraries. ACS Combinatorial Science 18, 438–443, doi:10.1021/acscombsci.6b00058 (2016).

