## Supplementary Information for "MRBLES 2.0: High-throughput generation of chemically functionalized spectrally and magnetically-encoded hydrogel beads using a simple single-layer microfluidic device"

**Supplementary Figures**

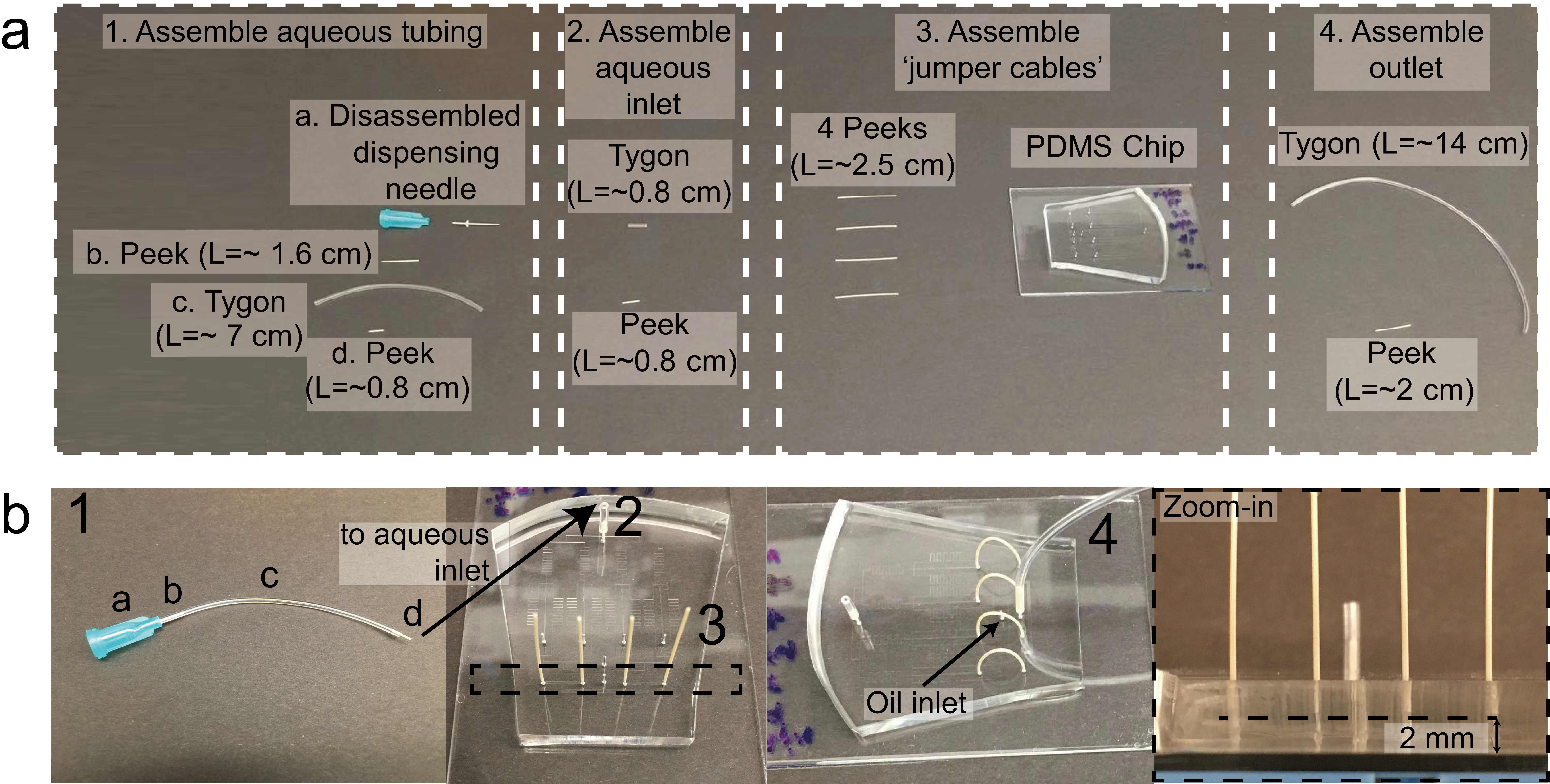

**Figure S1**. Assembly of single-layer high-throughput droplet generator using a ‘jumper cable’ strategy (see Supplementary Methods for step-by-step instructions). **(a)** Required tubing parts, including: (1) removable input assembly for aqueous phase (including (*a*) Luer fitting from disassembled blunt end Luer needle, (*b*) 1.6 cm Peek tubing, (*c*) Tygon tubing, and (*d*) 0.8 cm Peek tubing), (2) aqueous phase input assembly that remains connected to the device, (3) ‘jumper cables’ used to route droplets generated by individual flow focusers to a common device outlet, and (4) outlet tubing required to collect droplets from the device and route them to collection wells. **(b)** Assembly of individual tubing parts into the completed device: the removable aqueous phase input (1) is inserted into the aqueous phase input that remains connected to the device (2); ‘jumper cables’ are inserted into four outer ports in the droplet collection module (3), and then gently bent and inserted into to the corresponding outlets of four flow-focusers (4). The distance between the end of the PEEK tubing and the bottom of the PDMS device should be ~2 mm to ensure equal resistances and flow rates across the device and prevent device delamination (‘zoom-in’, right).

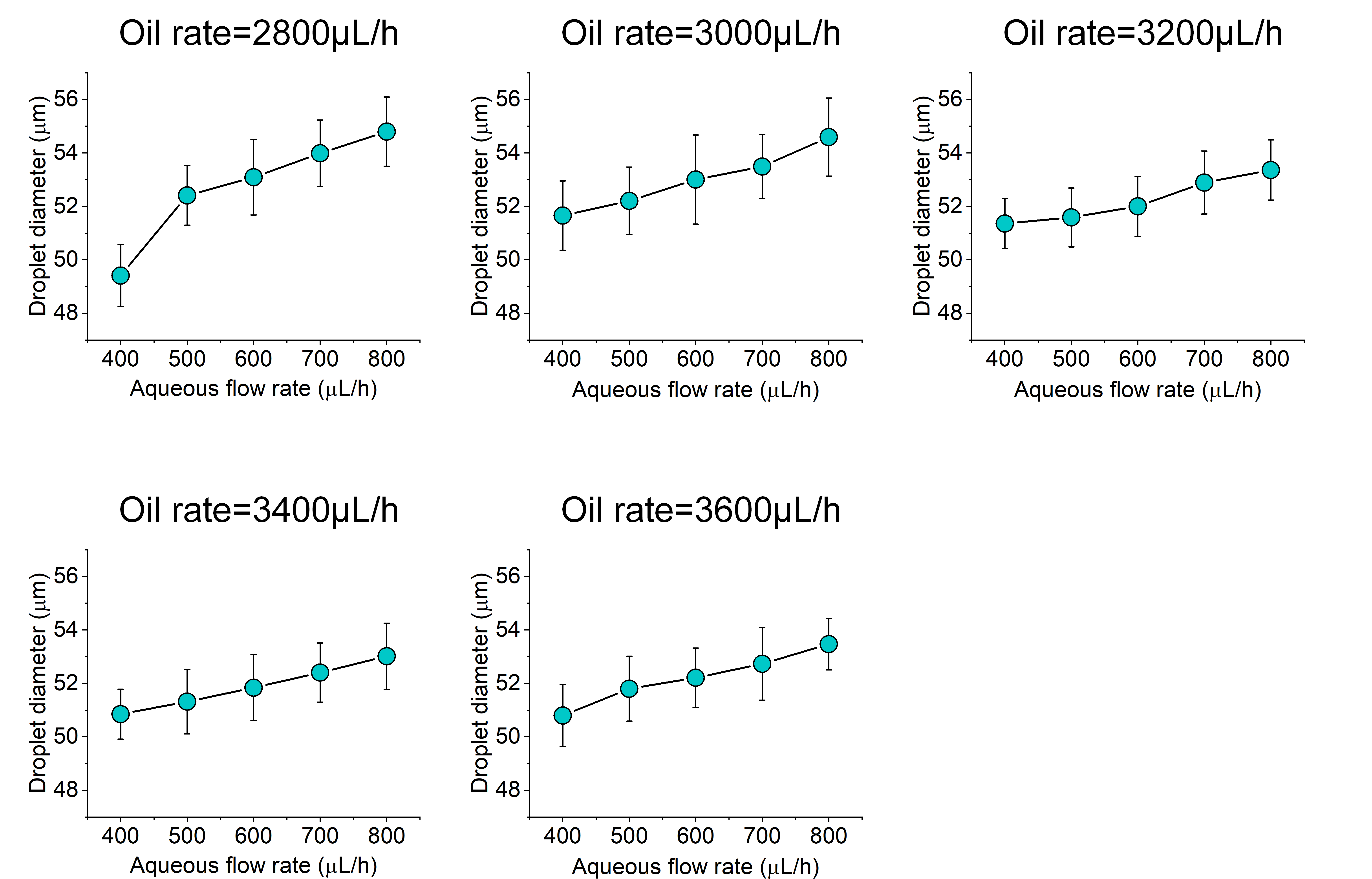

**Figure S2**. Measured droplet diameters as a function of aqueous and oil flow rates.

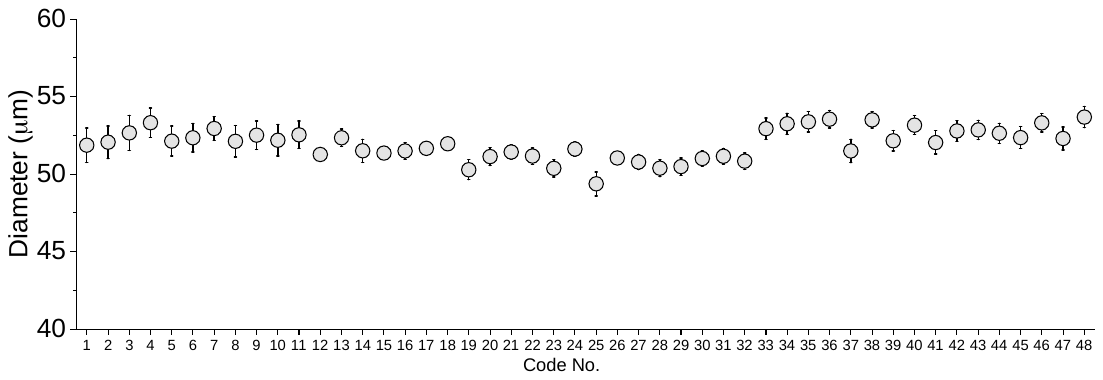

**Figure S3**. Measured bead diameters across all 48 spectral codes (mean ± standard deviation) after UV polymerization.

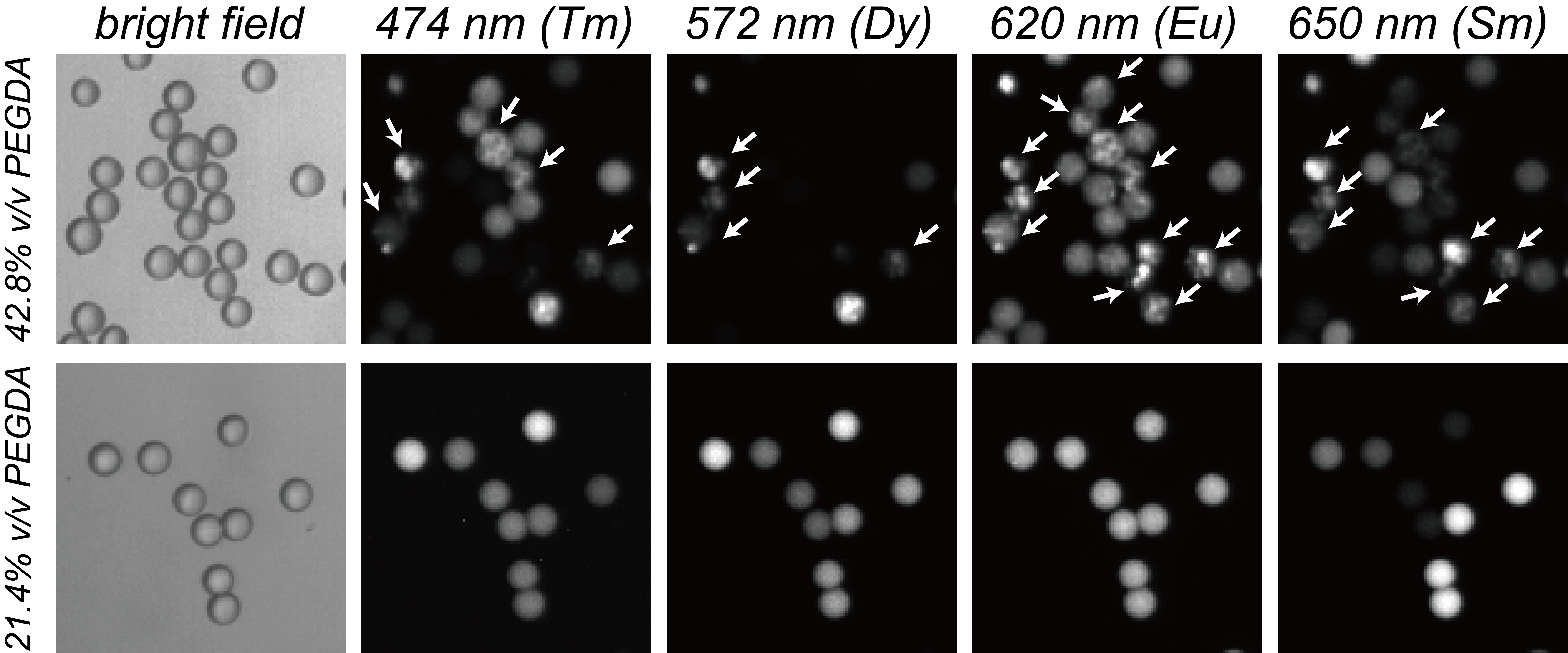

**Figure S4**. MRBLEs produced at 2 different PEG-DA concentrations imaged under bright field illumination and in 4 of the 9 lanthanide emission channels. High PEG-DA concentrations (42.8% v/v, top panels) lead to visible aggregation of Lns within the MRBLEs hydrogel matrix (visible as puncta within the right 4 lanthanide channels and highlighted by the white arrows); lower PEG-DA concentrations (21.4% v/v, bottom panels) significantly reduce aggregation and yield uniform intensities.

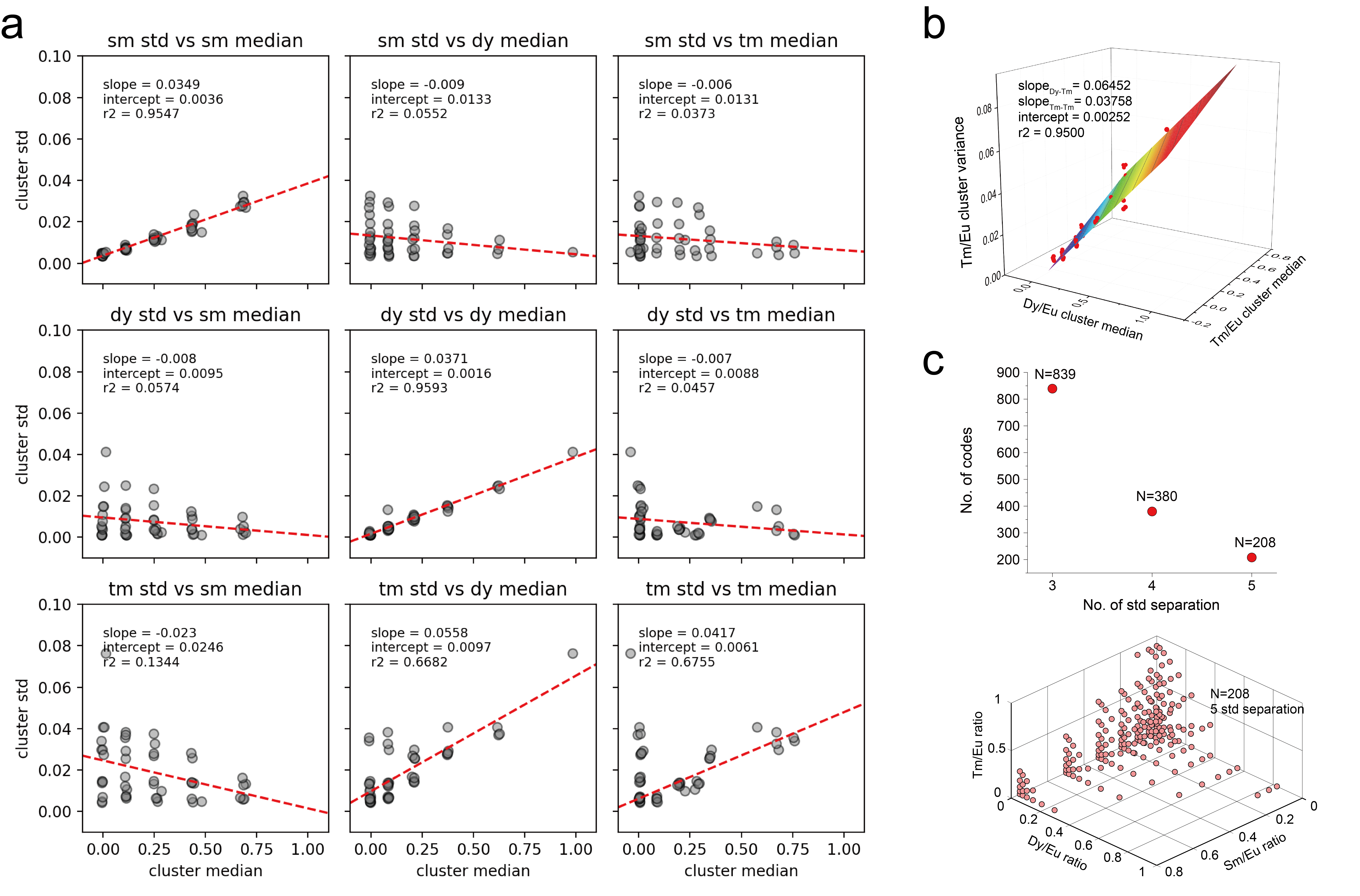

**Figure S5**. MRBLEs 2.0 calculated coding capacity. **(a)** Matrix of Sm/Eu (top), Dy/Eu (middle) and Tm/Eu (bottom) cluster variance as a function of cluster median. Note that the Tm/Eu cluster variance depends on the medians Dy/Eu and Tm/Eu values for each cluster. **(b)** Surface fitting of Tm/Eu cluster variance *vs.* cluster medians of both Dy/Eu and Tm/Eu. **(c)** Calculated code clusters yielding 839, 380 and 208 codes when requiring 3-, 4- and 5-standard deviation spacing between each code cluster. Reference ratios from the 5-standard deviation spacing code set (bottom) are well-separated and typically used by our laboratory for MRBLEs synthesis.

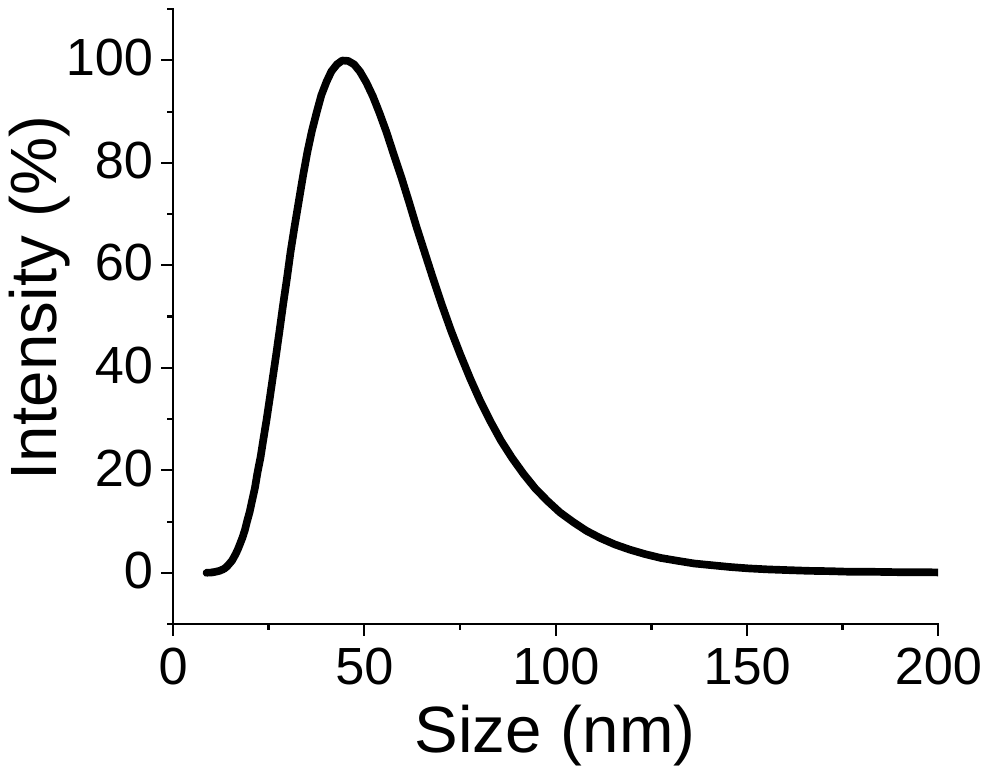

**Figure S6**. Size distribution for synthesized Fe_3_O_4_ magnetic nanoparticles as measured via dynamic light scattering (DLS) (mean = 46 ± 19.9 nm).

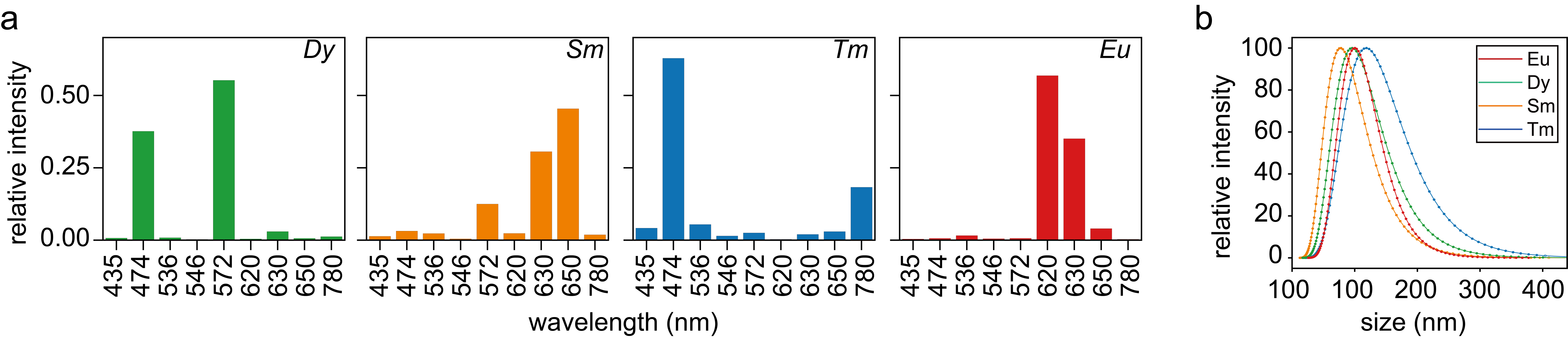

**Figure S7**. Emission spectra and size for synthesized Ln nanoparticles. **(a)** Normalized emission spectra for YVO4:Eu, YVO4:Sm, YVO4:Dy and YVO4:Tm Lns under excitation at 292 nm. **(b)** Size distribution of synthesized Ln nanoparticles as measured by DLS.

**Supplementary Table**

**Table S1**. 48 code reference table (volumetric ratios of master mixtures and target intensity ratios).

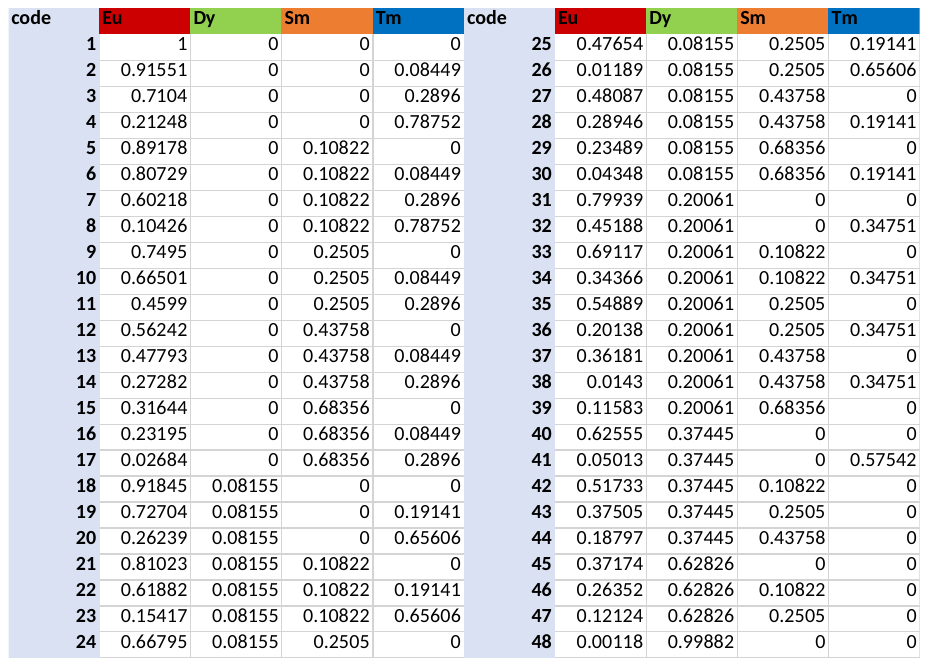

**Supplementary Methods**

*Chip assembly*. AutoCAD designs of all devices are provided as Supplemental Files and in an associated OSF repository (<https://osf.io/jvnpc/>). Each device requires 4 different sets of tubing (**Figure S1a**): (1) removable aqueous inlet tubing that can be connected to a syringe to load Ln/polymer mixtures and can be easily disconnected from the device and washed between codes, (2) aqueous inlet tubing that remains attached to the device and connects to this removable assembly, (3) 4 different ‘jumper cables’ that route droplets from individual production channels to a single collection outlet, and (4) outlet tubing that directs droplets to a multiwell plate or tube for collection.

Contact with metal can lead to premature polymerization of the Ln/polymer/photoinitiator and subsequent device channel clogging. To prevent this, we removed all metal from components required to introduce aqueous reagents into the device and route and collect droplets and prepared the necessary tubing as follows:

1. To assemble the removable aqueous inlet tubing (**Figure S1a,** 1), we first disassembled a blunt end needle with a Luer lock connection (McMaster-Carr, PART #75165A684, **Figure S1a,** 1*a*). Next, we cut two short pieces of PEEK tubing (L ≈ 1.6 cm and L ≈ 0.8 cm) (0.010” ID × 0.020” OD, ZEUS ,, **Figure S1a,** 1*b* and 1*d*) and one piece of Tygon tubing (L ≈ 7 cm) (0.125” ID ×0.25” OD, McMaster-Carr, PART #6516T14, **Figure S1a,** 1*c*). To create a final assembly, we inserted the two pieces of PEEK tubing in either end of the Tygon tubing and then inserted the longer piece of PEEK tubing into the Luer lock connection.
2. To assemble the aqueous inlet tubing that would remain connected to the device throughout the bead synthesis process (**Figure S1a,** 2), we cut additional 0.8 cm and 0.8 cm pieces of the same Tygon and PEEK tubing, respectively, and inserted the PEEK tubing into the Tygon tubing.
3. To prepare the tubing to be used for the ‘jumper cables’, we cut 4 additional 2.5 cm lengths of the same PEEK tubing (**Figure S1a,** 3).
4. To prepare the outlet collection tubing, we cut a 14 cm piece of Tygon tubing and a 2 cm piece of the PEEK tubing and inserted one into the other (**Figure S1a,** 4).

Next, we inserted the aqueous inlet tubing that would remain connected to the device into the aqueous inlet, the outlet collection tubing into the outlet (with the PEEK tubing inserted into punched device inlets in all cases), and all 4 jumper cables into the punched ports at the beginning of the collection chambers (**Figure S1b,** 1-3). Next, we gently bent the PEEK jumper cables and inserted them into the punched outlet ports at the end of the corresponding flow focuser channels (**Figure S1b,** 4). In all cases, PEEK tubing should be well inserted into the PDMS but leave an approximately 2 mm gap between the end of the tubing and the device slide to prevent delamination; tube ends should be aligned to an equal distance from the slide by eye to ensure equal resistance along each flow pathway.

*Droplet microfluidics set-up*. We visualized droplets using a very low-cost inverted bright field microscope (IQCrew 40X-200X Science Discovery Series Inverted Microscope, Amazon, $49.99) with three objectives (4X, 10X and 20X) coupled to a USB high-speed camera (Thorlabs, DCC1240M); to obtain a wider field of view, we replaced the 10X eyepiece with a C-mount 0.5X relay lens (AmScope). To inject aqueous and oil phases into the device at constant flow rates (volumetric flow), we used 2 syringe pumps (Pump 11 Elite Infusion Only Single Syringe, Harvard Apparatus). Prior to starting flow, we assembled the PDMS device by inserting the permanently-connected aqueous inlet tubing, ‘jumper cables’ and outlet collection tubing (**Figure S1b,** 4). To begin droplet production, we first introduced the oil phase at a flow rate of 3200 μL/h. After we saw a small drop of oil emerge from the aqueous inlet, we connected the removable aqueous phase tubing assembly containing the first Ln/polymer mixture and started the injection of the aqueous phase at a flow rate of 600 μL/h. For the first 1 minute of droplet production, we directed the outlet tubing to a ‘waste’ well (thereby providing time for droplet production to stabilize and reducing polydispersity) by clamping the tubing in place using a 3-way helper clamp ~ 4 cm above the top of the well. After 1 minute, we moved the plate to position the output tubing above a target well containing 80 μL of the oil phase solution (to inhibit the interaction between emulsion drop and the plastic bottom). Each 250 µL aqueous phase formulation was usually fully injected and all droplets formed within ~10 min; towards the end of droplet production, the droplets became progressively smaller (as visualized in the orifice region on the screen). At this point, we: (1) unplugged the aqueous phase and let the oil back-flow to the aqueous inlet to clear any remaining aqueous phase formulation from the channels, (2) moved the plate to direct the output to the ‘waste’ well (to collect any remaining droplets pushed by oil), and (3) disconnected the removable portion of the aqueous phase inlet tubing from the device, flushed it with DI water (to remove any remaining aqueous phase material), and dried it with air. At this point, we transitioned to the next code by refilling the syringe with the next Ln/polymer mixture and repeated droplet production.

*Lanthanide synthesis*. We synthesized lanthanide nanophosphors largely as described previously^1^. All chemical reagents and poly(acrylic acid) sodium (NaPAA), 45 wt% water solution for nanophosphor synthesis were purchased from Sigma-Aldrich (St. Louis, MO) and used without further purification. Microwave synthesis was performed using a Biotage Initiator (Biotage AB, Uppsala, Sweden). Synthesized nanophosphors were purified with DI water by ultrafiltration using Amicon Ultra-15 centrifugal filter units with a 30,000 MWCO (Millipore, Billerica, MA). The filtration was performed 4 times with ~40 mL DI water each time, resulting in a final 6 mL suspension slurry with a nanophosphor concentration of ∼50 mg/mL in water. Dynamic light scattering measurements established that produced Lns occupied a narrow size distribution (~100-nm in diameter, **Figure S7a**), with observed excitation spectra consistent with prior measurements (**Figure S7b**).

*Magnetic nanoparticle synthesis*. Magnetic nanoparticles were synthesized using co-precipitation and thermal decomposition methods. First, we prepared a mixture of 9 mL NaPAA (45 wt% water solution), 1.011 g KNO_3_ (Sigma-Aldrich) and 28.931 mL DI water in a reaction tube. Next, we incubated this reaction tube in a water bath preheated to 100 ^o^C and stirred at medium speed. We then added 6 mL of FeCl_2_∙4H_2_O (Sigma-Aldrich, 1 M in DI water, filtered) to the reaction. After the addition of this Fe^2+^ solution, we added 2.069 mL NH_3_∙H_2_O (Sigma-Aldrich, 30-33% NH_3_ in H_2_O) and observed the immediate formation of a black precipitate. We continued to heat the reaction at 100 ^o^C for 2 h with stirring, then transferred the slurry to a 50-mL falcon tube, and centrifuged at 4000 rpm for 10 min. We removed the supernatant after this spin step and purified it by ultrafiltration using Amicon Ultra-15 centrifugal filter units with a 30,000 MWCO to wash away any remaining NaPAA by DI water. The filtration process was identical to the lanthanide nanophosphors except that we resuspended the final product in ~3 mL DI water to yield a concentration of ∼150 mg/mL.

*Dynamic light scattering (DLS)*. We measured the size distribution of all of the synthesized lanthanide nanophosphors and magnetic nanoparticles using a DLS instrument (Brookhaven Instrument Nanobrook Omni). Prior to DLS, we diluted the Ln solution 40X and filled 1/3 of the cube; for the magnetic nanoparticles, we diluted them by 100x and again filled only 1/3 of the cube.

**Reference**

1.	Nguyen, H. Q.; Baxter, B. C.; Brower, K.; Diaz-Botia, C. A.; DeRisi, J. L.; Fordyce, P. M.; Thorn, K. S., Programmable Microfluidic Synthesis of Over One Thousand Uniquely Identifiable Spectral Codes. *Advanced Optical Materials* **2017,** *5* (3), 1600548.
